# Extracting neuronal activity signals from microscopy recordings of contractile tissue: a cell tracking approach using B-spline Explicit Active Surfaces (BEAS)

**DOI:** 10.1101/2020.12.15.422837

**Authors:** Youcef Kazwiny, João Pedrosa, Zhiqing Zhang, Werend Boesmans, Jan D’hooge, Pieter Vanden Berghe

## Abstract

Ca^2+^ imaging is a widely used microscopy technique to simultaneously study cellular activity in multiple cells. The desired information consists of cell-specific time series of pixel intensity values, in which the fluorescence intensity represents cellular activity. For static scenes, cellular signal extraction is straightforward, however multiple analysis challenges are present in recordings of contractile tissues, like those of the enteric nervous system (ENS). This layer of critical neurons, embedded within the muscle layers of the gut wall, shows optical overlap between neighboring neurons, intensity changes due to cell activity, and constant movement. These challenges reduce the applicability of classical segmentation techniques and traditional stack alignment and regions-of-interest (ROIs) selection workflows. Therefore, a signal extraction method capable of dealing with moving cells and is insensitive to large intensity changes in consecutive frames is needed.

Here we propose a b-spline active contour method to delineate and track neuronal cell bodies based on local and global energy terms. We develop both a single as well as a double-contour approach. The latter takes advantage of the appearance of GCaMP expressing cells, and tracks the nucleus’ boundaries together with the cytoplasmic contour, providing a stable delineation of neighboring, overlapping cells despite movement and intensity changes. The tracked contours can also serve as landmarks to relocate additional and manually-selected ROIs. This improves the total yield of efficacious cell tracking and allows signal extraction from other cell compartments like neuronal processes. Compared to manual delineation and other segmentation methods, the proposed method can track cells during large tissue deformations and high-intensity changes such as during neuronal firing events, while preserving the shape of the extracted Ca^2+^ signal. The analysis package represents a significant improvement to available Ca^2+^ imaging analysis workflows for ENS recordings and other systems where movement challenges traditional Ca^2+^ signal extraction workflows.

## Introduction

In order to understand how complex cellular systems operate and interact with each other, it is essential to be able to record activity from many individual cells simultaneously. Fluorescent calcium (Ca^2+^) imaging, either with small organic Ca^2+^ indicators or with genetically encoded Ca^2+^ indicators (GECI), (1, 2) is a widely used method to study large amounts of cells simultaneously and examine their network activity. Since cytosolic Ca^2+^ changes are tightly linked to action potential firing (and thus activity) in excitable cells like neurons, this imaging technique allows inferring neuronal activity of a large cellular population in both the central and peripheral nervous systems (3). Recent improvements in Ca^2+^ indicator quality (higher quantum efficiency and therefore better signal to noise) and imaging technologies allow monitoring larger populations of neurons at higher spatiotemporal resolution.

An extra complexity with live imaging of cells is that they may not be stationary in the microscopic field of view, either because they traffic themselves or the tissue, in which they are embedded, is contractile. Recordings in the central nervous system and acute brain slices can be assumed to have static scenes where the only movements present are motion artifacts such as drift, as in brain slices, or cyclic movements, as induced by breathing in intravital recordings. However, recording activity from tissues with a predominantly contractile function, such as the heart or the intestine, or from *in vivo* imaging of awake animals (zebrafish, C. *elegans, etc*) presents unique challenges due to the drastically high level of movement caused by muscle contractions.

In the intestine, all motor activity is controlled by a continuous network of neurons and glia cells embedded in between two concentric muscle layers. This enteric nervous system (ENS) regulates gut functions such as motility, secretion, and absorption (4,5). To understand how the complex circuits in the ENS operate to produce functional output, it is necessary to record and analyze the activity of large populations of ENS cells.

A traditional analysis workflow in Ca^2+^ imaging starts with image registration of the recorded frames to correct for motion artifacts and slight underlying movements aiming to attain a completely static scene where each pixel represents the same physical location throughout all frames (6). This step, if successful, is followed by signal extraction, where the different cells of interest are delineated and their pixel intensity profiles are extracted. For the large majority of Ca^2+^ imaging experiments, this workflow is sufficient to efficiently analyze cellular activity profiles and has been used extensively in ENS Ca^2+^ imaging provided that contractions are restrained either pharmacologically, physically, or in combination (7,8).

Multiple different software packages have been developed to automate the signal extraction process and efficiently analyze the ever-longer recordings and ever-increasing Ca^2+^ imaging datasets (9,10). However, these automated analysis workflows also rely on an image registration step and assume that all objects in the image are spatially static after this step, in order to extract their signals. Contractile movements, as those in the intestine, can include complex deformations that cannot be compensated with rigid registration techniques. More advanced non-rigid registration techniques, which offer registration with a high degree of freedom to accommodate more complex deformations, can be used but they are susceptible to high noise levels and artifacts, two regularly occurring problems in Ca^2+^ imaging. The tight packing of neurons in small groups (ganglia), with their apparent overlap in optical recordings (Fig. 1A), is a first challenge that eliminates the use of classic segmentation workflows. Moreover, the rapidly oscillating fluorescence of active neurons in Ca^2+^ imaging (Fig. 1B) has a negative impact on the success rate of registration algorithms as these rely on pixel intensity or image feature matching and thus have endogenous problems with changes in intensity (7,11,12). ENS Ca^2+^ imaging combines the aforementioned challenges and thus urges the development of an alternative analysis workflow to delineate and track individual cells in moving tissues, and extract their signals throughout the recordings.

**Figure 1:**
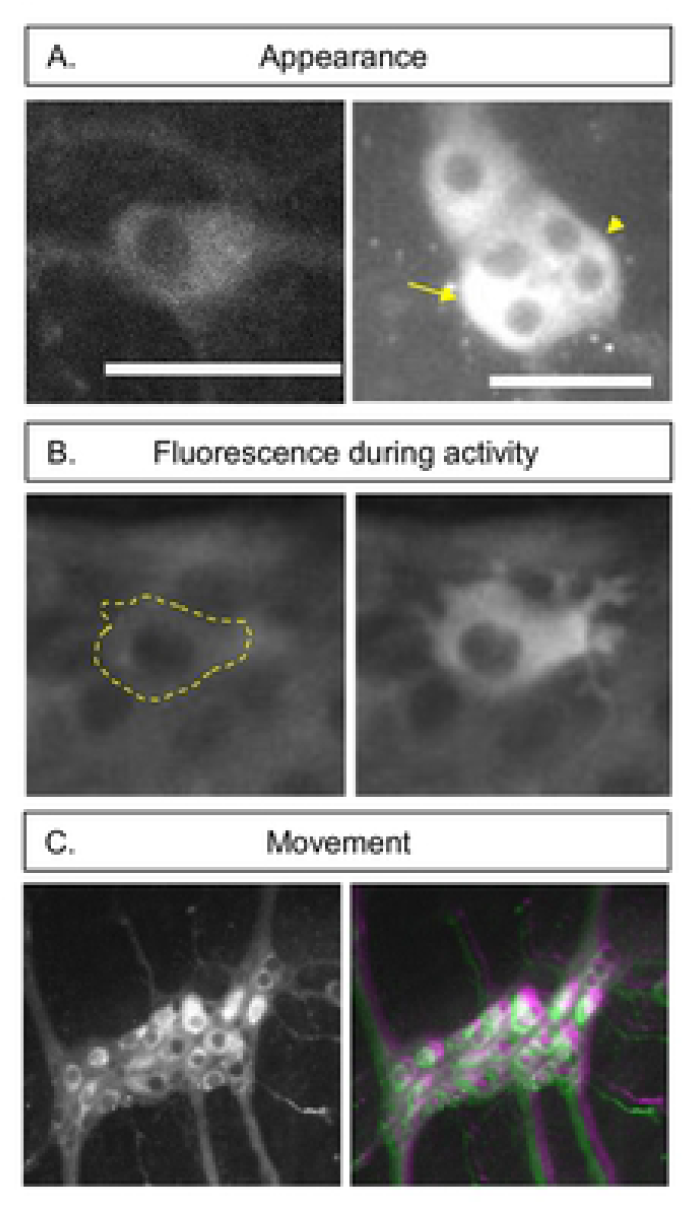
General features of Ca^2+^ imaging in the ENS. **A)** The appearance of an individual GCaMP expressing enteric neuron when not surrounded by other neurons (Left). The overlapping appearance of enteric neurons (arrow) and lack of clear borders (arrowhead) (Right). Scale bar represents 50 μm. **B)** an example of the fluorescence signal increases between a neuron at rest (left, and marked with a dashed line) and during activity (right). **C)** ENS ganglion (left) containing approximately 20 neurons. Imposed images of different timepoint (colorcoded in green and magenta) in an ENS Ca^2+^ recording (1 sec. interval between frames). The mismatch in colors indicates the amount of movement that can be present between 2 frames.

A viable alternative to registration in these complex scenarios is cell tracking. While tracking techniques have been extensively used in cell migration analysis and lineage tree construction (13–15), the low level-based segmentation techniques (15,16) that are normally used in these applications perform poorly in ENS recordings since they are prone to noise, variability in the edge intensity due to overlap, and cannot adapt to the blinking cell appearance between different frames (17). The existing region-based tracking techniques are not sufficient to segment complex structures based on their texture information (18,19). Moreover, they are ineffective when dealing with nonhomogeneous and overlapping objects, such as cells with bright cytoplasm and dark nuclei (Fig. 1A) as is the case with the expression of the common Ca^2+^ indicator GCaMP. Only one report, by Hennig *et. al*. (20) was published, in which nucleus tracking of ENS neurons was used, by means of edge detection where dark nuclei were identified and segmented in each frame to extract fluorescent GCaMP signals from their surrounding pixels. Practically, manual region-of-interest (ROI) selection remains the most commonly used approach to analyze ENS recordings, at least for those in which motion can be easily corrected. Recordings that rigid registration cannot stabilize are routinely disregarded.

Due to its ease of application and flexibility in handling cell division, the main method used in the cell tracking field is segmentation, based on implicit functions such as level-sets (21–23). However, the large flexibility in this implicit topological representation can easily produce incorrect results (24) especially in low signal to noise ratio (SNR) recordings. In these situations, explicit functions such as explicit active contours (25) perform better as they depend on parameters and therefore their evolution is more restricted and faster to calculate (26). The main disadvantage of explicit active surfaces is the inability in handling cell division, which is not relevant in the specific context of tracking neurons (16). In this paper, we implement B-spline-Explicit Active Surfaces (BEAS) as developed by Barbosa *et. al*. (27) which allows the application of local and global region-based energy terms in segmentation, as originally developed for level-set segmentation (28), while controlling contour smoothness and keeping the computational cost low (27,29). This method is suitable to segment heterogeneous objects (such as cells with dark nuclei, with varying degrees of brightness and edge clarity, Fig 1B) and to apply multiple local and global energy terms to reach that goal.

In this paper, we use the BEAS framework on 2D microscopy recordings to track and analyze multiple cells within a contractile and moving ENS tissue. Apart from employing multiple global and local energy terms to direct contour evolution, we also use a competition penalty to limit and manage overlap between neighboring cells. Furthermore, we develop ‘double contour (DC)’ tracking, a novel method that couples the development of two contour layers and takes advantage of the typical appearance of GCaMP expressing cells. Due to the nuclear exclusion of GCaMP, these cells present in Ca^2+^ imaging recordings with a dark nucleus and a bright cytoplasm, the edges of which are respectively tracked by the two layers. This DC method enables accurate cell tracking even in the absence of visible external borders. We describe the elements in the Ca^2+^ imaging and cell tracking algorithm developed and make this information freely available for external use.

In conclusion, we aimed to develop a set of techniques to better extract cellular activity levels from Ca^2+^ imaging recordings of non-static moving cells (Fig. 2). To this end, we used the ENS as a model system harboring fairly complex movement and activity-dependent intensity changes. The resulting workflow is however flexible and can be used to analyze other cellular recordings by tweaking the contour parameters to match the specific application.

**Figure 2:**
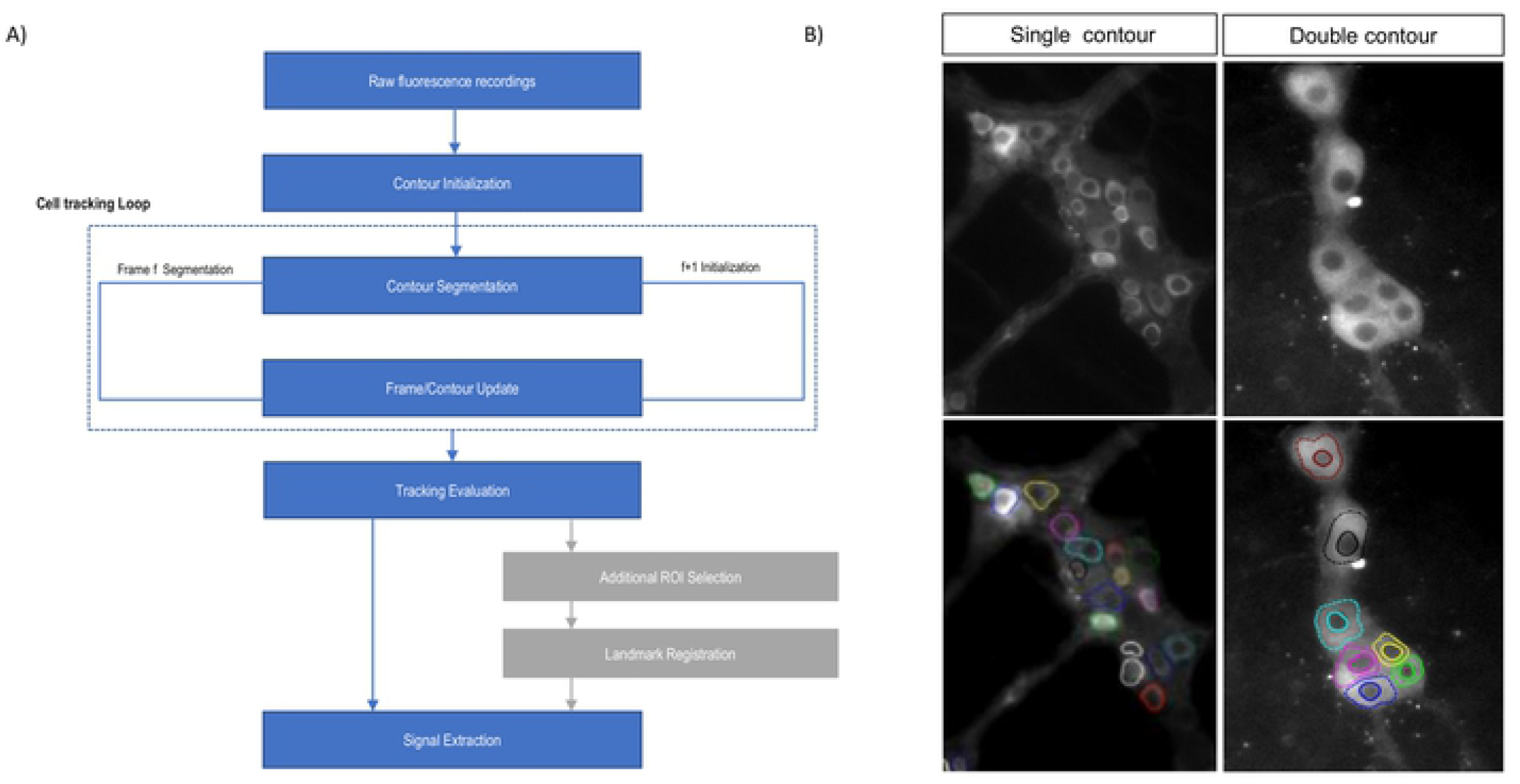
A) diagram of the processing workflow including the main steps of cell tracking (Blue) and the optional ROI tracking (grey). B) Example of the performance of the cell tracking procedure of multiple cells in ENS recordings, using one-layer tracking (left) and double contours (right)

## Methodology

The workflow for the proposed cell tracking approach starts by drawing an ellipse around the cell to initialize the contour. This step is followed by deforming the contour iteratively by applying forces on individual contour control points until the functional energy minimum is reached as an initial segmentation step, which theoretically overlays the contour with the cell’s boundary. The initialization is followed by the cell tracking loop, which consists of a series of consequent segmentation tasks on individual frames, where each contour in a frame is used to initialize the contour’s segmentation on the following frame. During an intermediate step, parametric information about the contour is calculated and the contour center is also recalculated to be in the geometric centroid of the produced contour shape to ensure that the new center is inside the cell in each next frame, even if there was movement between frames (Suppl. Fig. 1). By stringing the segmentation results together, we acquire both the location of individual cells as well as their contours throughout the entire recording (Fig. 2 B).

The goal of this approach is to use these dynamic contours as regions-of-interest (ROIs) from which the mean intensity signal is extracted to accurately represent Ca^2+^ activity of cells in a non-static setting. These contours are then evaluated by the user. Furthermore, the tracked cell locations can also be used as landmarks to optionally track or displace additional and manually created ROIs, in cases where a tracked cell’s contour was not satisfactory or when tracking additional ROIs posthoc is desired (Fig. 2 A).

### B-Spline Explicit Active contours algorithm (BEAS)

We implement the B-Spline Explicit Active Surfaces (BEAS) (30) framework developed and optimized for segmenting and tracking heart chambers in echocardiography (29–31). The method uses an explicit function to represent the boundary of an object, where coordinates of the contour points are explicitly given as a function of the remaining coordinates i.e., *x*_1_ = *ψ*(*x*_2_,…, *x_n_*) where *ψ* is defined as a linear combination of B-spline basis functions

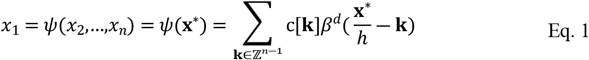

where *β^d^*(.) is the uniform symmetric B-spline of degree d. The knots of the B-splines are located on a rectangular grid, with a regular spacing given by *h*. The coefficients of the B-spline representation are gathered in c[k]. For this 2D segmentation problem, a polar coordinate system was chosen.

The evolution of the contour is governed by the minimization of the energy term E. This energy has two elements, the image data term E_*d*_ and an internal energy E_*r*_.

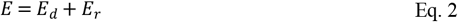

### Data attachment

#### One-layer Contour

The data attachment energy term can be defined, following the BEAS formulation, as:

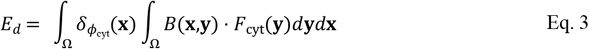

where *F*_cyt_(**y**) is the energy criterion driving the evolution of the contour and *B*(**x,y**) is a mask function in which the local parameters that drive the evolution are estimated. *δ*_*ϕ*_cyt__(**x**) is the Dirac operator applied to the level set function *ϕ*(**x**) = Γ(**x**\*) – *x*_1_ which is defined over the image domain Ω. The mask function B(**x,y**) for a node (neighborhood radius) is specified as a column of pixels of length ρ in the normal direction centered around a contour node. The value of ρ is chosen a priori, based on the expected margin (frontier) size between objects and the rate of movement between frames. When segmenting GCaMP expressing cells, it is logical to set this parameter to be slightly smaller than the approximate radius of cells, to avoid detecting the cytoplasm-nucleus edge instead of the intracellular interface. The degree of visibility of a cell’s border in Ca^2+^ imaging is quite variable as its strength is based on the Ca^2+^ concentration inside the cell of interest as well as that of adjacent cells. Moreover, the imaging conditions and imaging system chosen also impact the cell’s appearance (Fig. 1A). Therefore, we chose a flexible localized energy term introduced by Yezzi et al. (32) (Eq. 4), to maximize the difference of mean intensity inside and outside each contour node.

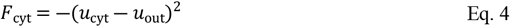

Where *u*_cyt_ and *u*_out_ are the mean intensity values in the cytosolic region (inside the cell) and the region outside of the cell, respectively.

#### Double Contour

In live fluorescent imaging (*eg*. in Ca^2+^ imaging), the interface between the bright cytoplasm, which can be dim if intracellular Ca^2+^ concentrations are low, and the heterogeneous background may lack contrast and as such limit cell tracking capability. GCaMP expressing cells have a bright cell body and a dark nucleus because the GCaMP molecule molecules do not enter the nucleus. Therefore, a second, and often sharper interface, between the dark nucleus and the bright cytoplasm emerges. This interface is stable and has a predictable (dark) inner side and (bright) outer side. Therefore, we developed a coupled two-layer active contour segmentation of cells. The two layers delineate the nucleus-cytoplasm and the cytoplasm-background interfaces, respectively. The inner layer *ϕ*_nuc_ is delineating the stable shape of the nucleus while the outer contour *ϕ*_cyt_ attempts to delineate cell outer borders forming a “double contour”. The image-data energy term can then be defined as:

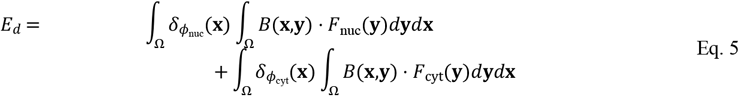

with *F*_nuc_ following *F*_cyt_ in its definition (Eq. 4). The double contour produces a more stable contour progression and keeps the contour attracted to cells in the event of non-visible cellular borders (Fig. 3). It also allows the extraction of the signal that originates from the cytoplasm pixels only, which improves the signal to noise ratios of the extracted mean fluorescence.

**Figure 3:**
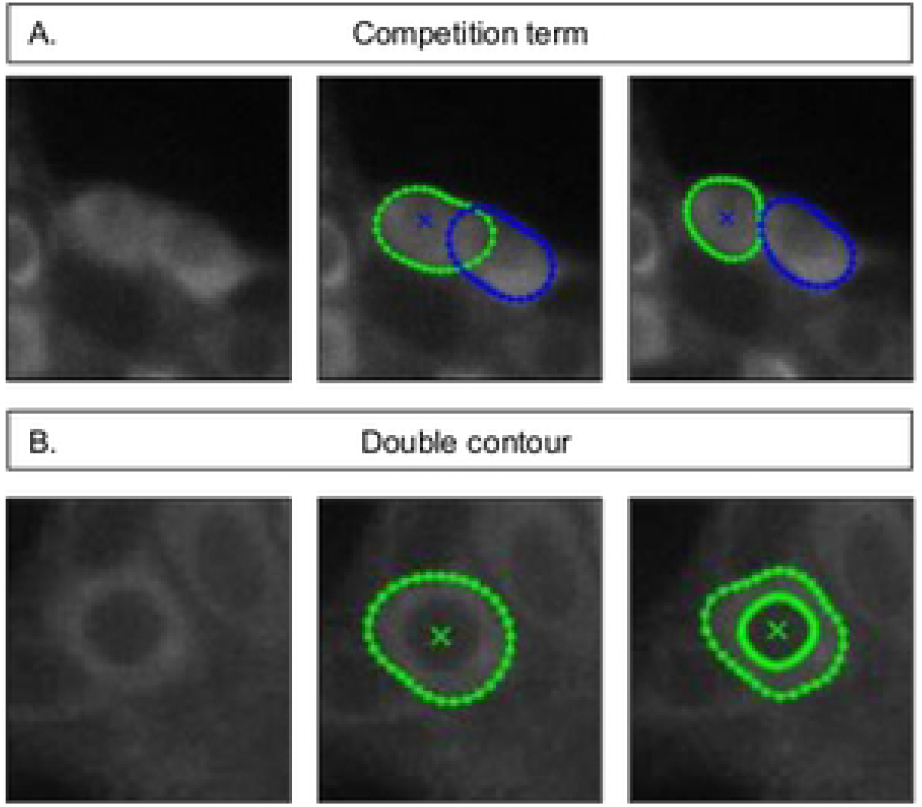
The effect of a competition term on neighboring contours (top) The contour of a cell using 1-layer vs double contour in a GCaMP expressing neuron (bottom)

### Data regularization

The energy term E_*r*_ relates to curvature, size, and size difference compared to the previous frame. We use prior knowledge about the properties of ENS neurons to impose local and global penalties to guide the contours and ensure that segmentation results and contour shapes will be plausible in their curvature, size, and size differential between timesteps. The regularization term E_r_ is defined as:

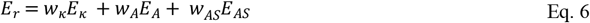

The curvature energy term E_κ_ limits the negative local mean curvature since cell bodies mostly have positive curvature. The local curvature gradient term is given by:

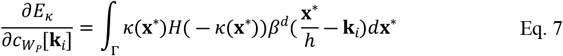

Where *κ* is the local mean curvature which is calculated efficiently as reported within the BEAS framework (29,33) and H is the Heaviside function.

The area energy term E_A_ keeps the size of the contour within a reasonable range, where A represents the area within the contour. The parameters A_min_ and A_Max_ ensure that the contour does not engulf bigger image regions. The equation for local energy calculation is governed by:

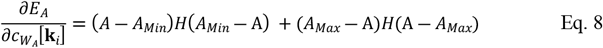

Next, we add the area stability energy term E_AS_, which is a global energy term that attempts to minimize the change of the area within the contour keeping its size in a reasonable range for a cell, since apparent size changes are not real but are due to intensity variations or edge contrast changes and not caused by actual cell size changes.

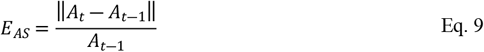

The weights *w_κ_, w_A_* and *w_AS_* in Eq. 6 are chosen by the user based on image dimensions and cell types.

### Contour Competition

It is common for cells in microscopy recordings to appear overlapping, as an image is a projection of all fluorescent elements in the focus of the objective lens. Especially in widefield microscopy recordings where images result from many different in- and out-of-focus planes (34). This effect is minimized in confocal and multiphoton excitation approaches, but optical overlap remains an issue due to limited optical resolution. While banning overlap completely can facilitate interpretation of the extracted data, it does not represent the scene correctly and can lead to tracking errors. Therefore, we impose a competition penalty that allows a slight contour overlap to account for the optical overlapping effect while preventing contours from jumping between cells or engulfing multiple cells. We opted to impose a proximity penalty between neighboring contour nodes, as implemented previously in BEAS (35), to limit contour expansion into neighboring contours and reduce overlap (*E^dist^*),

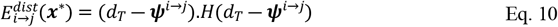

where *d_thresh_* represents the minimal distance parameter, *ψ* is a signed distance map between each node of the contour i against all nodes of contour j (and vice-versa), and *H* is the Heaviside operator. Note that *H* equals one only in nodes with *ψ* lower than *d_thresh_* and zero in the remaining nodes. Therefore, it only applies penalties in the neighboring regions of the contours (35).

We also added a stronger penalty for actual overlap on both contours (*E_overlap_*) producing a cell competition effect controlled by the cell competition weight parameter *w_c_* (Fig. 3) that is a *priori* chosen.

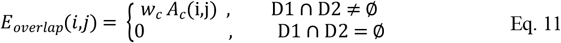

With D1, D2 being the pixels belonging to contour i and j, respectively and *A_c_* is the area of overlap between two cells. Then Eq. 6 for contour i with a neighboring contour j can be rewritten to include the competition terms:

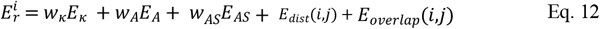

### Landmark-based geometrical transformation and ROI tracking

While we aim at effective cell tracking in every scenario using BEAS cell tracking, there are known challenges that can constrain tracking using active contour methods. These challenges include parameter sensitivity causing the algorithm to be suboptimal for some cells in the field of view, despite being successful for other cells. Therefore, we introduce a robust optional step that uses the tracked locations of *N* cell contours using the BEAS approach as landmarks to find the optimal geometrical transformation T that represents the movement in the recorded scene between frames. The optimal parameters *θ*^*^ of T are estimated by minimizing the similarity measure d (36, 37) which represents the Euclidian distance of the cell contour coordinates between two frames:

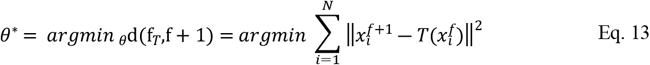

With 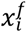 being the centroid coordinates *x* of the contour *i* in frame *f*. This geometrical transformation allows us to move additional ROIs selected manually by the user *posthoc* throughout the recording frames by performing the geometrical transformation T (38) on the positions of the ROIs.

### Implementation details

Initialization is done by manually selecting ellipses that roughly overlap with the targeted cell bodies. These ellipses are fed as initial contours to the first frame segmentation step. The result of contour segmentation in a frame is then used for contour initialization in the next frame. Practically, the neighborhood radius ρ determines the range of cell movement between frames that is detectable by the segmentation step. ρ is chosen empirically to detect large movements without extending far off the cell edge and losing its ability in finding local cell edges and relies on multiple parameters including image resolution and relative movement (Suppl. data). During the initialization step, overlap was not allowed to simplify the initial contour interactions and limit entanglement in later segmentation steps. We choose to represent the B-spline contours in polar coordinates because cell bodies appear as closed ellipses. Therefore, the geometry functions took the form of r=ψ(θ). The geometrical center of the contour shape is calculated and the pole of the contour coordinates translated to this point after each time step (Suppl. Fig. 1). This step is essential as contours cannot be represented as a polar curve if the pole (coordinates’ origin) is outside of the cell contour.

The angular discretization factor denoting contour boundary nodes was set empirically to 32 nodes with regular angular interval dθ. When applied to the experimental recordings, this setting was found to provide a good balance between shape flexibility and representation at a reasonably low computational cost. We measured its effect and that of other parameters in a dedicated parameter sensitivity test. New contour nodes were resampled after the translation step to preserve the accuracy levels of the discretization and maintain the regular interval dθ. This was done by using linear interpolation of the contour nodes’ coordinates (r’, θ’) for polar angles θ’ with a regular dθ interval. A modified gradient descent with feedback step-adjustment was used to perform the energy criterion minimization as explained in previous BEAS implementations (11,30). Runtime was linearly dependent on the number of cells and the image size. The geometrical transformations T used in landmark-based ROI tracking is implemented in the form of a polynomial affine transformation (39).

## Results

### A. Segmentation strategy evaluation

To objectively evaluate the presented segmentation strategies, we created an artificial dataset that simulates the Ca^2+^ imaging scenes, featuring movement at rates similar to what is measured in ENS recordings, several intensity-change patterns that represent Ca^2+^ activity, overlapping neighboring cells with similar baseline intensities, and multiple blurred frames to mimic out-of-focus imaging frames (Fig. 4).

**Figure 4:**
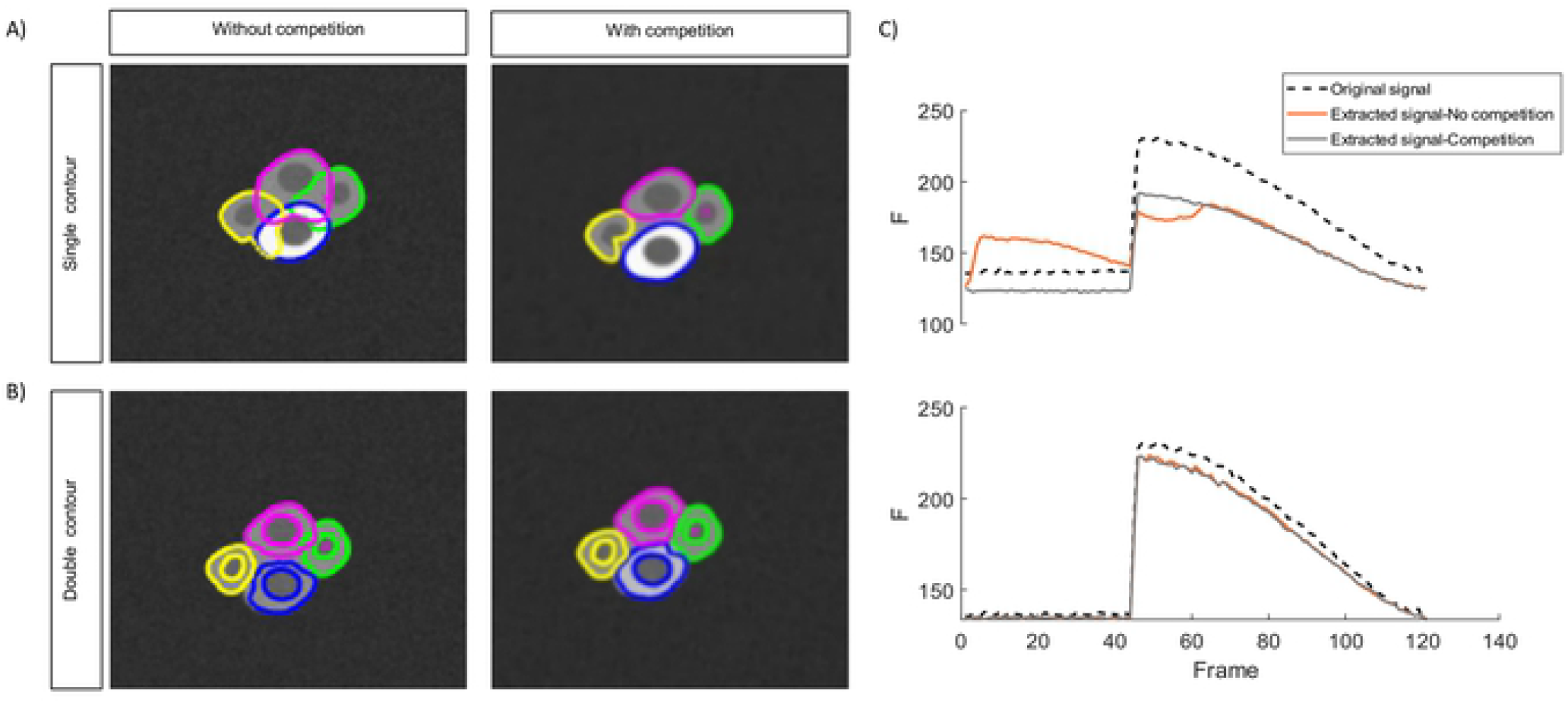
Tracking of overlapping cells with the same base intensity level during rest and different intensity levels during activity using **A)** one-layer contours without a competition parameter (left) and with a competition penalty added (right) and **B)** double contours without a competition parameter (left) and with a competition penalty added (right) **C)** Extracted signals from a cell using one-layer contours (top) and double contours (bottom)

We analyzed the signals in this dataset using four different approaches and compared how the extracted signals matched the ground truth signals. Using one-layer segmentation which targets the cytoplasmic border only, without the competition term, expectedly yields poor results, with contours overlapping significantly as the contour nodes cannot find clear edges or intensity gradients (Fig. 4A, left). As a result, the extracted signals are contaminated with information from neighboring cells. On the other hand, using the competition term in addition to the fixed global curvature term anchors the contours and restricts their shapes to prevent them from taking over neighboring cells (Fig. 4 A, right), which improves the extracted signal quality drastically. In contrast, the double contour segmentation maintains the general shape even without a competition penalty due to the coupling between the two segmentation layers although a slight overlap can be observed. The small overlaps, in this case, are alleviated when the competition term is added (Fig. 4 B).

Signals extracted from the artificial dataset confirm that one-layer contour tracking, without competition, is not reliable in extracting the original signal. This is shown in Fig. 4 C top, where the activity from the neighboring cell appears in the activity trace of the measured cell (Fig. 4 C, top row, red trace). Double contour segmentation and one-layer contours with competition terms, have no such issues and allow extraction of an accurate signal shape. This is especially the case for double contour segmentation, where the raw fluorescence level is closer to the original because now the dark nucleus pixels can be excluded from the calculation of the cytoplasm intensity (Fig. 4 C, bottom).

### B. Parameter sensitivity analysis in an artificial dataset

The impact of each of the selected parameters on segmentation and tracking results using active contours in both one-layer and the double contour methods are shown in Fig. 5. The first parameter ρ, the neighborhood radius (Fig 5A), expectedly has, for smaller values, a big influence on the tracking results.

**Figure 5:**
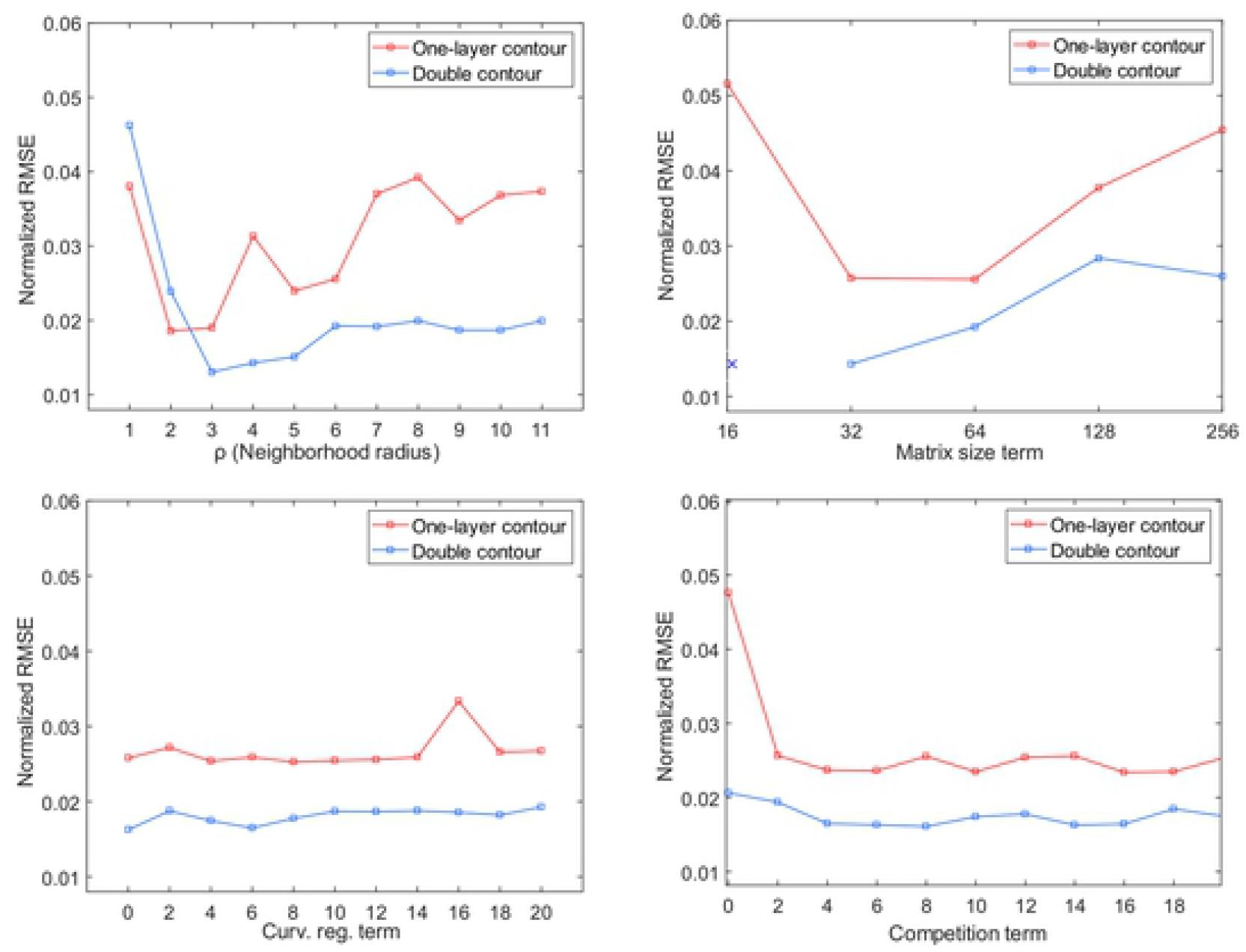
Parameter sensitivity analysis: comparison between one-layer vs. double contours where the Y-axis is the normalized RMSE of the extracted signals compared to the raw signals: (top left) effect of radius length values on segmentation sensitivity, double contour segmentation has less segmentation error for all values in the relevant range > 3. (top right) effect of the number of contour nodes: higher RMSE for one-layer segmentation for all values, note the segmentation failure of double contour method at low (e.g. sixteen) contour nodes number, as indicated with x. Curve regulation term (bottom left). Competition term (bottom right): lack of competition term causes high normalized-RMSE for one-layer segmentation.

Afterwards, the tracking is stable for several values until the radius is too large and tends to encounter multiple edges simultaneously. The second parameter, the matrix size, which determines the number of discretized contour nodes negatively affects the tracking at smaller matrix sizes (fewer number of nodes) for both one- and double contour tracking, with failure to track cells in case of double contour segmentation with the lowest (only sixteen) number of nodes (Fig. 5B). This indicates that due to its extra complexity, double contour segmentation is more sensitive to the number of nodes. While the curvature term does not affect the accuracy of segmentation (Fig. 5C), the addition of a competition term does improve segmentation, especially for the one-layer segmentation option. The effect of a competition term with double contour segmentation is negligible in this dataset (Fig. 5 D).

### C. Experimental results

When applied to actual recordings, we find that the proposed approach successfully tracks cells throughout significant tissue movements (Fig. 6, Top panel), allowing us to reliably resolve Ca^2+^ peaks from the extracted signals (Fig. 6, bottom panel).

**Figure 6:**
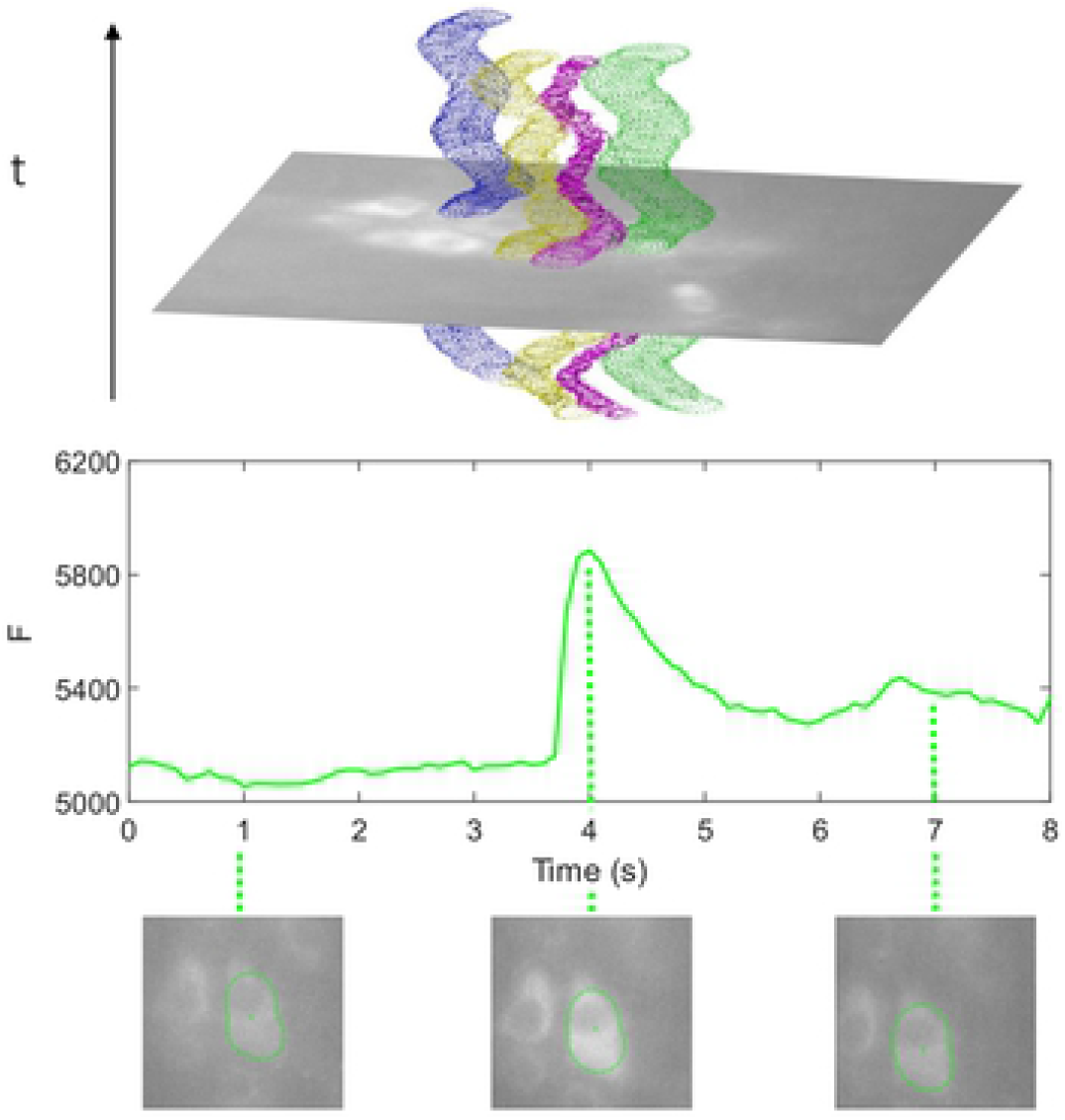
Contours of multiple cells and cell movement (top). Pixel intensity signal from the one tracked cell 2+ in the top panel and the contour appearance at multiple time points before, during and after a peak in Ca^2+^ activity.

To compare with traditional analysis methods, we analyzed recordings with both the new contour tracking method as well as with manual routines, involving motion correction and rectangular ROI selection by a blinded expert. For that purpose, we used datasets of 3 recordings to compare the degree of similarity of the signals extracted by the traditional method against the one-layer contour and double contour methods, respectively (Fig. 7 A, B). We found that Ca^2+^ profiles are very comparable in shape between extraction from tracked cells versus manually drawn ROIs, with a normalized root-mean-square error (RMSE) of 0.093 and 0.114 for one-layer or double contours respectively (Fig. 7 B).

**Figure 7:**
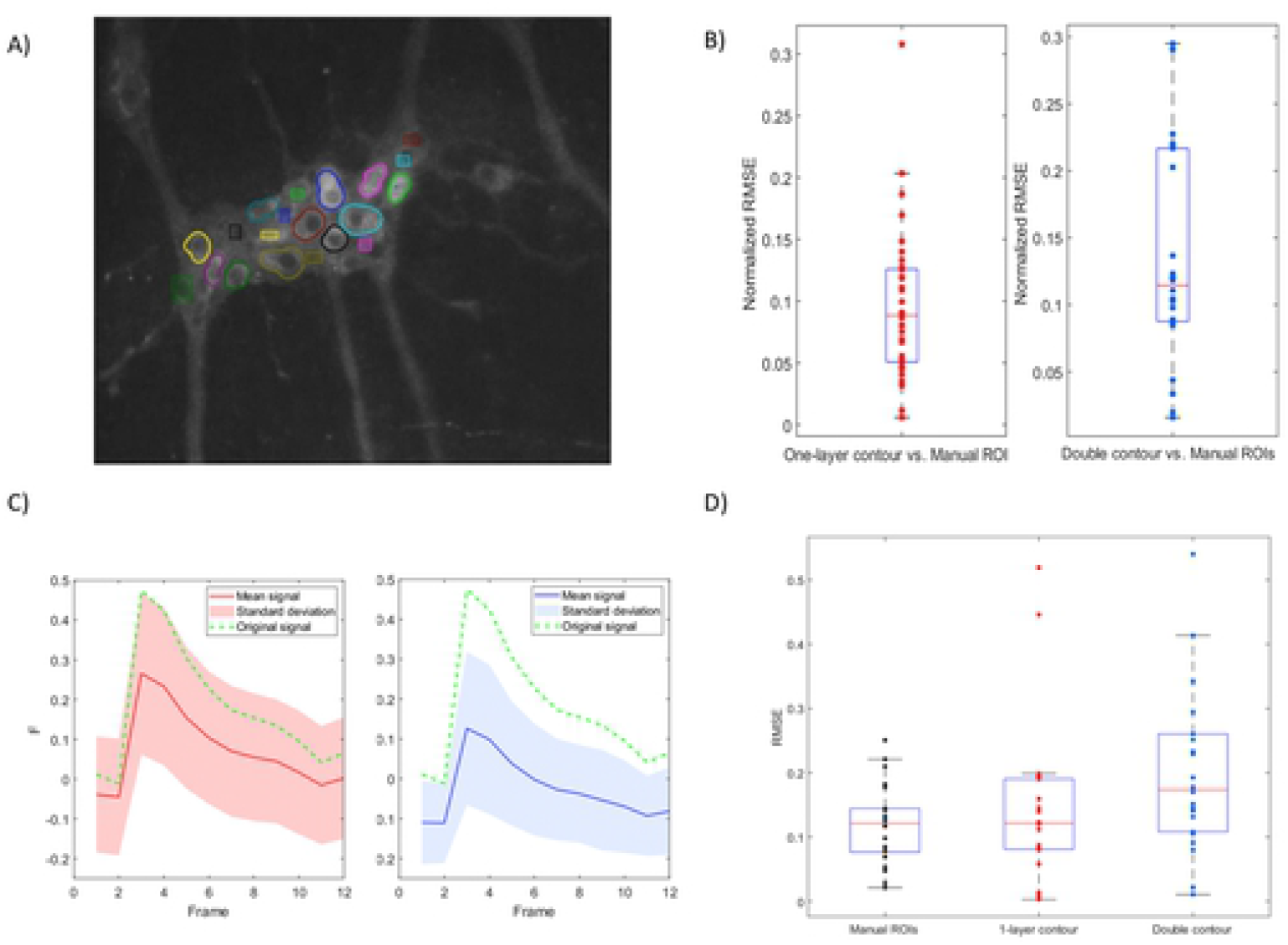
A) Contours of tracked cells and manual rectangular ROIs which are moved based on the contours tracked. B) RMSE of signals extracted using 1 layer (red) and double contour (blue) versus manually selected ROIs by an expert. C) Comparison of the extracted signals (red/blue mean, light red/blue standard deviation) against a ground truth artificial peak (dotted green) using 1-layer contour tracking (left) and double contours (right) based on 22 cells. D) RMSE of signals extracted from 22 cells injected with the artificial peak using manual ROIs (left), 1 layer (center) and double contour (right) versus the ground truth.

True validation of our analytical approach is not straightforward as it requires assessing the quality of signal extraction against a ground truth signal. Since the latter should be fully known, yet embedded in a context that holds all the biological, optical, and experimental complexity, we generated artificial Ca^2+^ peaks in real Ca^2+^ recordings in the cytoplasm of multiple moving cells (Suppl. movie 1). Then we used the contour cell tracking method to re-extract the Ca^2+^ signals and compared the extracted signals to the original planted signal. We found that the Ca^2+^ peak shape was preserved in most cells (Figure 7 C), and the median RMSE in the one-layer and double contour approach to be 0.1216 and 0.1738 (based on 22 tracked cells) to be comparable to that of the manually selected ROIs with an RMSE of 0.1212 (Fig. 7 D). The higher RMSE values of DC tracking are simply due to a few complete tracking failures, see *Discussion*.

Finally, we used the newly developed tracking approach on actual recordings of ENS tissue (Suppl. movie 2, 3) including movement in x and y and out-of-focus frames. We found that with optimized parameters one-layer segmentation, proves reliable to track many cells in the field of view. Notably, the segmentation procedure performs well despite the presence of blurry out-of-focus frames (Fig. 7), which is an important advantage compared to edge-based segmentation techniques (40). In double contour segmentation, we observed less overlap of contours without a competition penalty resulting in good reliability in cells with non-visible edges. However, this method had more difficulties in scenes with faster movement and was expectedly not robust in cells without contrast between the nucleus and cytoplasm. While this new approach performs well, it is unavoidable that cell tracking fails to resolve some cells with challenging appearance or location in the image. The developed landmark ROI-tracking exploiting the known trajectory of successfully-tracked cells proved to be a useful and robust tool to overcome this challenge with minimal computational power needed (Suppl. movie 4). In addition, it gives the researcher the additional ability to extract signals from smaller structures, like cell processes or glial cells (Fig. 7 A).

## Discussion

Given the complexity of Ca^2+^ imaging in the contractile ENS tissues, where a scene not only contains moving cells but these cells also display irregular fluorescence intensity changes (6), traditional methods based on image registration and ROI selection are cumbersome and prone to failure during signal extraction and quantitative analysis (3, 41–43). Additionally, low-level cell tracking techniques cannot function reliably in this scenario due to multiple reasons, including low signal-to-noise ratio (often the case in live imaging), cellular overlap and variable cellular edges, which depend both on the imaging system and the labeling approach, as well as on the activity of the cell (44). In this paper, we developed a cell tracking algorithm targeted specifically to track neurons in such a challenging contractile scenario, with the additional complexity that cells in Ca^2+^ imaging have blurry borders and constantly change fluorescence intensity. Our method successfully tracks blinking cells in moving ENS tissue, without the need for non-rigid image registration. The extracted temporal signals are comparable in quality to manual, expert-selected ROIs. Furthermore, the tracked cell coordinates allow additional rectangular ROI tracking and add robustness and flexibility to the workflow to process the most challenging recordings.

### A. Comparison of segmentation strategies

In an artificial dataset that was created to simulate cell shape and behavior, specifically having moving cells without clear borders, we found the competition term to be important in one-layer cell tracking as the contours overlap and the contour nodes do not find edges to adhere to in their absence. The addition of a competition term and a significant curvature term prevents them from taking over neighboring cells resulting in good signal extraction.

The performance of cell segmentation in this simulated dataset is consistently improved by using the novel double contour method. The double contour uses the inner nucleus contours as a natural anchor that restricts the outer contour from taking over neighboring cells. It can conserve the shape with low, or even without, competition and curvature terms. However, the advantage of the coupled double contour approach is limited in recordings with large cell displacement between frames as it depends on tracking the smaller nucleus from its position in the previous frame. In this case, higher image acquisition rates are required, which adds complexity to the imaging setup, generates bigger datasets, and causes longer processing times. Nevertheless, we consider the double contour approach to be powerful in its application to GCaMP based recordings, as the reporter is genetically prevented from entering the nucleus, leaving the nucleus dark and thus enabling accurate tracking of cells and signal extraction selectively from the cytoplasm.

### B. Parameter sensitivity analysis

Active contours are heavily reliant on multiple parameters and can be sensitive to parameter values limiting their robustness (45). We quantified the effects of the global penalty terms on the algorithm’s performance in both the one-layer and double contour strategies, by extracting and comparing the signals from simulated data. We observed general similarity in sensitivity to the studied parameters, except for the inability to track cells when using only a few contour nodes in the double contour, which is at odds with the increased complexity of this strategy. The introduction of the cell competition term improved cell tracking when using one-layer contours, reduced the error rate to similar values as obtained by double contours in this dataset. Although the curvature term did not increase the accuracy of the extracted signal in the simulated dataset, it plays an important stabilizing role to the cell contours in real recordings, especially in blurred images or in frames where segmentation is struggling to delineate cells returning to baseline fluorescence. We found that tracking is generally insensitive to a wide range of parameter values in the simulated dataset despite our efforts to introduce the most challenging conditions, which all together indicate that the performance of the algorithm is robust.

In general, the global penalty terms are valuable to limit segmentation failure, which is a drawback of active contour segmentation (28). However, they do not show significant effects on tracking results of cells that are already well within the means of the method, as shown in Fig. 5.

### C. Experimental results

As the aim of the new approach was to extract accurate Ca^2+^ signals from experimental data, we compared contour tracking to the traditional extraction method and found a high similarity of the extracted signals between the two methods. We used an artificially embedded Ca^2+^ peak to measure the similarity to the ground truth and found that these planted peaks were indeed detected in most cells, demonstrating the applicability of the contour tracking workflow. The artificially embedded Ca^2+^ peaks were then used to compare the quality of the signal extraction using the two contour types against the traditional extraction method. Results from the one-layer contours were highly similar to those of the traditional method in their error between the extracted signal and the ground truth values of the artificial peak (Fig. 7). We observed slightly lower average similarity between the ground truth signal and double contour method, which was mainly due to instances where the method failed to track those neurons without contrast between the nucleus and cytoplasm, which we, in order to be as close to reality as possible, also included in the dataset. This is easily mitigated by using the additional ROI tracking option, which we introduced to extract signals from cells for which contour tracking is inaccurate (Fig. 2A).

Practically, we find one-layer tracking to be robust in recordings with blurry out-of-focus frames and its stability largely depends on the neighborhood radius ρ in relation to movement intensity in-between frames. Furthermore, the introduction of a cell competition term improves cell tracking and reduces the error in experimental recordings. Double contour tracking on the other hand is useful when the recording is not blurry and the movement in-between frames is generally less than the nucleus diameter. The latter limits the applicability in recordings with substantial displacement due to rapid muscle contractions, especially when fast image acquisition is not feasible. Its main advantage, which results from the inner nucleus contour acting as an anchor to the outer cellular contour, is the ability to track overlapping cells without clearly visible borders, a common sight for ENS neurons in the submucosal layer (4). The landmark-based ROI tracking possibility for manually-added ROIs provides a useful addition that allows tracking challenging cells, which the active contours method fails to correctly segment. It is a useful tool as it does not require re-running the tracking workflow and is applied post-hoc, providing a robust option fully controlled by the user.

## Conclusion

To satisfy the need for a robust analysis tool for Ca^2+^ imaging in moving and contractile tissues, we introduced an efficient hybrid approach to track cell bodies relying on local region-based terms in evolving the contour, avoiding the disadvantages of region-based segmentation (Fig. 2). We further developed a novel ‘double contour’ or coupled-layers tracking algorithm that takes advantage of the fact that cells in genetically encoded Ca^2+^ imaging techniques appear with dark nuclei. We quantified the method’s performance in an artificial dataset that simulates experimental challenges under different parameter values and compared the two tracking algorithms. We then tested the algorithm’s robustness in tracking neurons in various ENS tissue Ca^2+^ recordings and demonstrate, using embedded artificial Ca^2+^ spikes, that the method reliably captures these spikes and represents them in the extracted signals. We expanded the analysis possibilities by implementing land-mark based ROI tracking, which increases the robustness of the workflow for challenging datasets. Finally, we packaged the workflow as a MATLAB GUI to enable efficient analysis of Ca^2+^ imaging datasets with a non-static scenery. The technique can be used on other cellular recordings by tweaking the contour parameters to match the specific application.

## Acknowledgements

The authors thank Tobie Martens for manual cell delineation and ROI selection on Ca^2+^ recordings. Animal tissues used in this study were taken from mice house and euthanised according to the guidelines and procedures as approved by the Animal Ethics committee of KULeuven. The authors’ work is supported by the Research Foundation Flanders (FWO) grant G.0929.15, G.OH1816N and I001918N and Hercules AKUL/11/37 and AKUL/15/37 (to P.V.B.).

## Supplementary information

see separate file

## Supplementary movies

**Movie 1**: The artificial dataset used for parameter sensitivity analysis: 7 overlapping cells with bright cytoplasm and dark nuclei representing moving and overlapping neurons in a noisy and blurry scene.

**Movie 2**: Example of the one-layer contours approach to track multiple neurons in ENS tissue

**Movie 3**: Example of the double contours approach to track overlapping neurons, with similar intensity baselevels in ENS tissue during stimulation.

**Movie 4**: Example of the one-layer contours approach in combination with ROI tracking to track multiple neurons during large deformation in ENS tissue.

